# Sleep deprivation exhibits age-dependent effect on infra-slow global brain activity

**DOI:** 10.1101/2025.09.12.675828

**Authors:** Yibing Yang, Weiwei Zhao, Yutong Mao, Tianxin Mao, Yao Deng, Yifan Yang, Hengyi Rao, Xiao Liu

**Affiliations:** Department of Biomedical Engineering, The Pennsylvania State University, University Park, PA, 16802, USA; Center for Functional Neuroimaging, Department of Neurology, University of Pennsylvania, Philadelphia, PA, USA; Chronobiology and Sleep Institute, Department of Psychiatry, University of Pennsylvania, Philadelphia, PA, USA; Institute for Computational and Data Sciences, The Pennsylvania State University, University Park, PA, 16802, USA

## Abstract

Infra-slow (< 0.1 Hz) global brain activity, quantified by the global mean BOLD (gBOLD) signal in resting-state fMRI, is elevated during sleep and closely associated with cerebrospinal fluid (CSF) dynamics, a key mechanism for the brain waste clearance implicated in neurodegenerative disorders such as Alzheimer’s disease (AD). However, the effect of sleep deprivation on gBOLD activity and its interaction with aging remain poorly understood. Using a rigorously controlled in-laboratory total sleep deprivation (TSD) protocol, we demonstrate that TSD significantly increases both the amplitude of the gBOLD signal and its coupling with CSF flow, suggesting a compensatory mechanism that may enhance glymphatic clearance following acute sleep loss. Notably, these TSD-induced enhancements exhibit robust age dependency, with diminished responses in individuals at midlife (40-50 yrs). The absence of this compensatory mechanism in these midlife participants may exacerbate age-related impairments in neurotoxic clearance and increase dementia susceptibility, thereby offering mechanistic insights into the nexus between sleep disruption, aging, and neurodegeneration.

## Introduction

Sleep is essential for maintaining homeostasis and restoring optimal brain functions^1,2^, potentially including the clearance of neurotoxic proteins^3^. Recent studies have consistently suggested the important role of a glymphatic system in brain waste clearance, which exhibits enhanced activity during sleep to presumably facilitate the removal of toxic proteins^3–5^. Consistent with this notion, sleep loss impairs the clearance function, reducing the brain’s capacity to eliminate toxic proteins and leading to their accumulation^6–9^. For example, animal studies have demonstrated that sleep deprivation elevates levels of Aβ and tau, the key pathological markers of Alzheimer’s disease (AD), in both the interstitial fluid (ISF) and cerebrospinal fluid (CSF)^8,9^. In parallel, human studies indicated that even a single night of total sleep deprivation (TSD) was associated with greater retention of contrast agents and a significant increase in Aβ burden in critical regions such as the hippocampus and thalamus^6,7^.

Recent work has revealed an infra-slow (<0.1 Hz) global brain activity that is linked to sleep-dependent neurotoxic clearance processes^10,11^. This activity, manifested as pronounced peaks in the global mean blood-oxygenation-level-dependent (gBOLD) signal^12,13^, exhibits highly structured spatiotemporal patterns in both human and animal studies^13–15^. In human fMRI and monkey electrocorticography (ECoG) recordings, it propagates as waves across cortical hierarchies, while in mouse neuronal recordings, it appears as cascades of sequential activation^14,15^. Notably, these widespread cortical activations are accompanied by deactivations in subcortical arousal-regulating centers, consistent with associated fluctuations in arousal states^13,14^. Most importantly, this global activity is coupled to CSF flow, suggesting a potential mechanistic role in brain waste clearance^10^. Supporting this notion, the strength of gBOLD-CSF coupling has been associated with various AD-related pathologies, including the accumulation of Aβ and tau^16,17^. Topological changes of gBOLD activity has even been linked to the spreading pattern of Aβ in the early stages of AD^18^.

While the global BOLD signal displays elevated amplitude during sleep^19,20^, the effect of sleep deprivation on gBOLD activity and its coupling with CSF dynamics remain poorly characterized. Emerging evidence implicates age as a critical modulator of sleep-dependent waste clearance^21^, potentially influencing susceptibility to sleep deprivation-induced changes in gBOLD activity. A contrast-enhanced MRI study in clinical cohorts identified age-dependent declines in glymphatic clearance efficiency^21^. Normative aging populations exhibit a nonlinear trajectory of gBOLD-CSF coupling marked by accelerated deterioration beginning around the mid-50s^22^. These changes align with the well-documented age-related decline in sleep quality^23^, characterized by reductions in slow-wave sleep, increased fragmentation, and diminished overall sleep efficiency. Thus, the interaction between age and sleep deprivation on gBOLD activity may be critical for understanding the age-related decline in waste clearance, which may in turn underlie the heightened risk of dementia in midlife (40-50 yrs) adults^24,25^.

In this study, we leverage resting-state fMRI data from a rigorously controlled in-laboratory sleep deprivation experiment^26^ to investigate changes in gBOLD activity and its coupling with CSF dynamics after one night of TSD and following two nights of recovery sleep across different age groups. Our results demonstrate that TSD induced a significant increase in gBOLD signal amplitude, which returned to baseline levels after two nights of recovery sleep. Interestingly, the TSD effect is highly age-dependent and almost absent in midlife subjects aged 40-50 years. Furthermore, the impact of TSD was more pronounced in higher-order brain regions, particularly within the default mode network (DMN), compared with lower-order sensory-motor regions. Given recent evidence implicating the key role of gBOLD activity and gBOLD-CSF coupling in brain waste clearance^18,27^, the increased global brain activity after sleep loss may represent a compensatory clearance mechanism. However, this response appears to be impaired in midlife, potentially contributing to the age-related decline in waste clearance efficiency and increased risk of dementia during aging.

## Results

### TSD increases gBOLD activity and its coupling with CSF dynamics

Resting-state fMRI data were collected from 67 subjects (34.1 ± 8.9 years, 29 females) who participated in a 5-day, 4-night TSD experiment, with scans conducted in the mornings of Days 2 (Scan1), 3 (Scan2), and 5 (Scan3) (**Fig. 1A**). gBOLD activity was quantified using two metrics: the standard deviation of gBOLD signal to assess its fluctuation amplitude, and gBOLD-CSF coupling (see Methods for details) to measure its synchronization with CSF flow measured at the bottom of the cerebellum (**Fig. 1B-1D**). The TSD group (*N* = 51, 33.7 ± 8.8 years, 21 females) showed a substantial increase (i.e., more negative) in sex-adjusted (applied to all gBOLD metrics) gBOLD-CSF coupling during the TSD scan on Day3 (*p* = 0.0071, *paired t-test*) compared to the baseline scan (BS) on Day1, and the coupling returned to baseline level during the recovery scan (RS) on Day5 after two nights of recovery sleep (*p* = 0.00041, *paired t-test*). In contrast, the control group (*N* = 16, 35.4 ± 9.5 years, 8 females) showed no significant differences in gBOLD-CSF coupling between the TSD and BS scans (*p* = 0.78, *paired t-test*) or between the RS and TSD scans (*p* = 0.88, *paired t-test*) (**Fig. 1E**). Similar changes (*TSD vs. BS, p = 0*.*00014, paired t-test; RS vs. TSD, p = 0*.*0015, paired t-test*) are also observed for the gBOLD amplitude (**Fig. 1F**).

**Figure 1.**
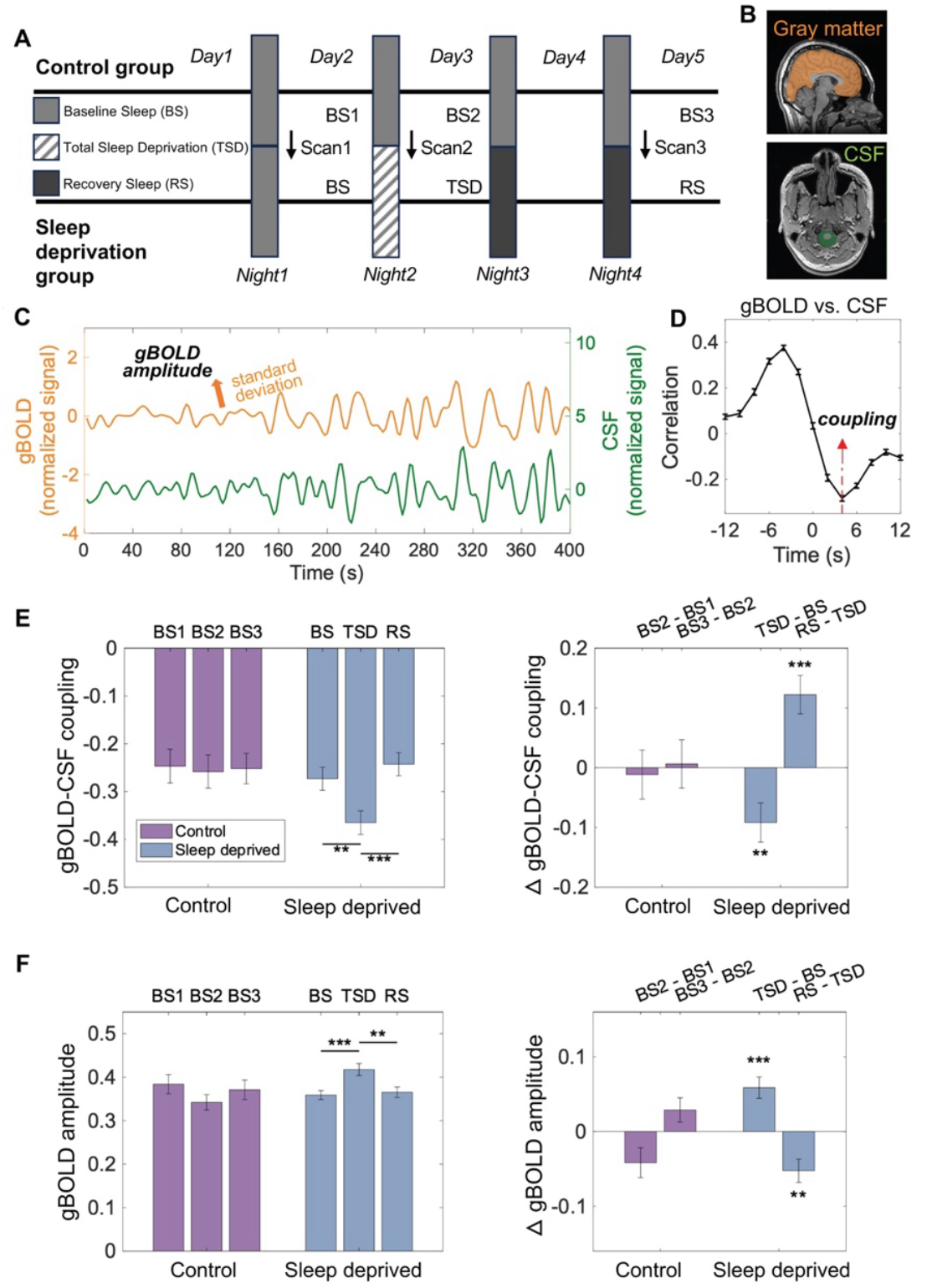
Sleep deprivation increased the amplitude of gBOLD activity and its coupling with CSF flow. **(A)** Illustration of the design of the 5-day, 4-night sleep deprivation experiment protocol. The sleep deprivation group (N = 51) completed four nights: Night 1 as baseline sleep (BS), Night 2 as total sleep deprivation (TSD), and Nights 3–4 as recovery sleep (RS). The control group (N = 16) underwent four consecutive nights of BS without TSD. **(B)** Example of gray matter mask to extract the gBOLD signal (the top panel) and CSF mask to extract the CSF signal from the bottom slice (the bottom panel). **(C)** gBOLD and CSF signals from a representative subject, with gBOLD signal averaged across all gray matter regions. The standard deviation of the gBOLD signal was computed to quantify its amplitude. **(D)** The cross-correlation function of gBOLD and CSF signals averaged across all subjects. The strength of gBOLD-CSF coupling was quantified by the correlation at the negative peak with +4 seconds time lag (red dashed line). (**E)** gBOLD-CSF coupling (left) and its changes (right) across three scans in sleep-deprived and control groups. Subjects who underwent total sleep deprivation showed a significant increase in gBOLD-CSF coupling during the TSD scan compared to the BS scan (p = 0.0071, paired t-test), and the coupling returned to baseline during the RS scan (p = 0.00041, paired t-test). No significant changes were observed in the control group (BS2 vs. BS1, p = 0.78; BS3 vs. BS2, p = 0.88, paired t-test). **(F)** The gBOLD amplitude (left) and its changes (right) over the three scans in sleep-deprived and control groups. Sleep-deprived subjects showed a significant increase in gBOLD amplitude during the TSD scan compared to the BS scan (TSD vs. BS, p = 0.00014, paired t-test), with amplitude returning to baseline during the RS scan (RS vs. TSD, p = 0.0015, paired t-test). Error bars represent the standard error of the mean (SEM) across subjects. Asterisks represent the significance level (*: 0.01 < p < 0.05; **: 0.001 < p < 0.01; ***p < 0.001)

### TSD effects on gBOLD activity are absent in midlife adults

To investigate whether the TSD-induced gBOLD changes are age-dependent, we divided the TSD subjects into three age groups: young (*N* = 20, age: 20-30 years, 8 females), early-midlife (*N* = 14, age: 30-40 years, 7 females), and midlife (*N* = 17, age: 40-50 years, 6 females) adults and compared their gBOLD metrics. The gBOLD-CSF coupling increased significantly after TSD in both the young (TSD vs. BS, *p* = 0.019, *paired t-test*) and early-midlife (TSD vs. BS, *p* = 0.0085, *paired t-test*) groups and then returned to baseline after recovery sleep (RS vs. TSD, *p* = 0.013 for young and *p* = 0.0037 for early-midlife, *paired t-test*). In contrast, no significant TSD-dependent changes were observed in the midlife group, with no differences between the TSD and BS scans (*p* = 0.82, *paired t-test*), or between the TSD and RS scans (*p* = 0.44, *paired t-test*) (**Fig. 2A**). Consistent with these observations, there was a significant interaction effect between age and scan (*p* = 0.037, *two-way ANOVA*) on gBOLD-CSF coupling.

**Figure 2.**
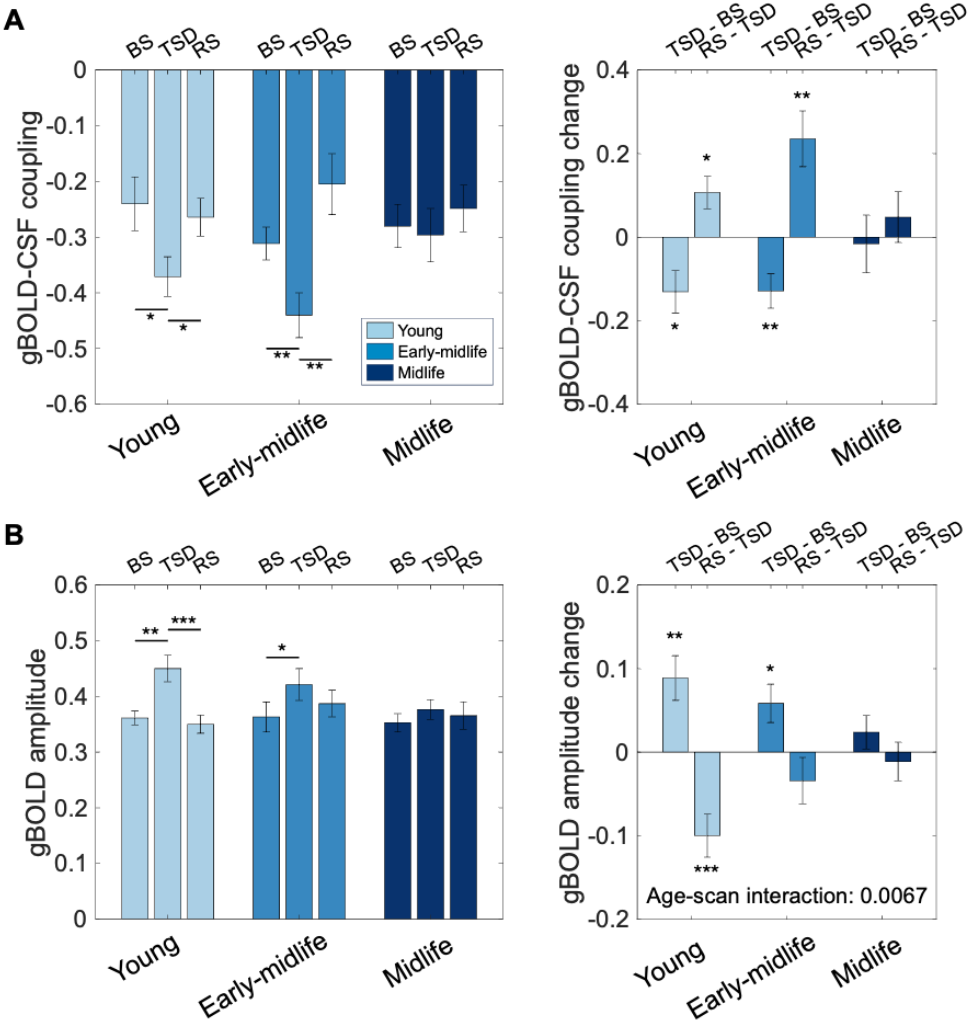
TSD effects on gBOLD activity across different age groups. (**A**) Changes in gBOLD-CSF coupling are significant in both young (TSD vs. BS, p = 0.0019; RS vs. TSD, p = 0.013, paired t-test) and early-midlife (TSD vs. BS, p = 0.0085; RS vs. TSD, p = 0.0037, paired t-test) groups, but not in the midlife group (TSD vs. BS, p = 0.82; RS vs. TSD, p = 0.44, paired t-test). There is a significant interaction effect between age and scan on coupling changes (p = 0.037, two-way ANOVA). **(B)** gBOLD amplitude increases after sleep deprivation in both the young and early-midlife groups (TSD vs. BS, p = 0.0036 for young and p = 0.025 for early-midlife, paired t-test) but not in the midlife group (TSD vs. BS, p = 0.25, paired t-test). Age and scan show a significant interaction effect on changes in gBOLD amplitude (p = 0.0067, two-way ANOVA). Error bars represent the standard error of the mean (SEM) across subjects.

A same analysis of gBOLD amplitude yielded similar results (**Fig. 2B**). In the young adult group, gBOLD amplitude increased significantly after TSD and then returned to baseline after recovery sleep (TSD vs. BS: *p* = 0.0036; RS vs. TSD: *p* = 0.00099, *paired t-test*). Similar but smaller changes were observed in the early-midlife group (TSD vs. BS, *p* = 0.025; RS vs. TSD, *p* = 0.24, *paired t-test*), whereas no significant changes were found in the midlife adults (TSD vs. BS, *p* = 0.25; RS vs. TSD, *p* = 0.64, *paired t-test*). The age-scan interaction was more significant (*p* = 0.0067, *two-way ANOVA*) than for gBOLD-CSF coupling. The head motion quantified by mean framewise displacement (FD) showed no significant correlations with either gBOLD-CSF coupling (*rho* = 0.13, *p* = 0.067, *Spearman’s correlation*) or gBOLD amplitude (*rho* = 0.0037, *p* = 0.96, *Spearman’s correlation*). The main findings remained robust and unchanged after controlling for head motion (**Fig. S1 and Fig.S2)**.

### TSD effects on gBOLD activity are pronounced in higher-order brain regions

Given that gBOLD activity shows more prominent reduction in higher-order DMN regions at the early stages of AD, which has been linked to early Aβ accumulation^18^, we examined whether TSD-induced gBOLD changes varied across the cortex and were biased toward specific regions. We quantified the topology of gBOLD activity using gBOLD presence, defined as the correlation between regional BOLD signals and the gBOLD signal^18,28^. Similar to gBOLD amplitude, gBOLD presence exhibited age-specific alterations in response to TSD, with minimal changes observed in the midlife group (**Fig. 3A**). In the young and early-midlife groups, TSD-induced increases in gBOLD presence were spatially heterogeneous, with larger effects observed in higher-order brain regions, including the bilateral posterior cingulate cortex (PCC), bilateral caudal anterior cingulate cortex (ACC), and bilateral superior frontal gyrus (SFG), which are all components of the DMN. These qualitative observations were supported by quantitative comparisons between the higher-order regions (DMN and frontoparietal network (FPN)) and lower-order regions of sensory-motor areas defined previously^18^. The increase in gBOLD presence was significantly greater in higher-order brain regions than in lower-order brain regions for both young (*p* = 0.0014, *two-sample t-test*) and early-midlife (*p* = 0.029, *two-sample t-test*) groups (**Fig. 3B**). In addition, the age-scan interaction effect on gBOLD presence was more significant in higher-order regions (*p =0*.*0067, two-way ANOVA*) than in lower-order regions (*p* =0.045, *two-way ANOVA*). After two nights of recovery sleep, the gBOLD presence deceased back to baseline with a similar pattern of spatial heterogeneity in the young group *(p* =0.0020, *two-sample t-test*) (**Fig. S3**).

**Figure 3.**
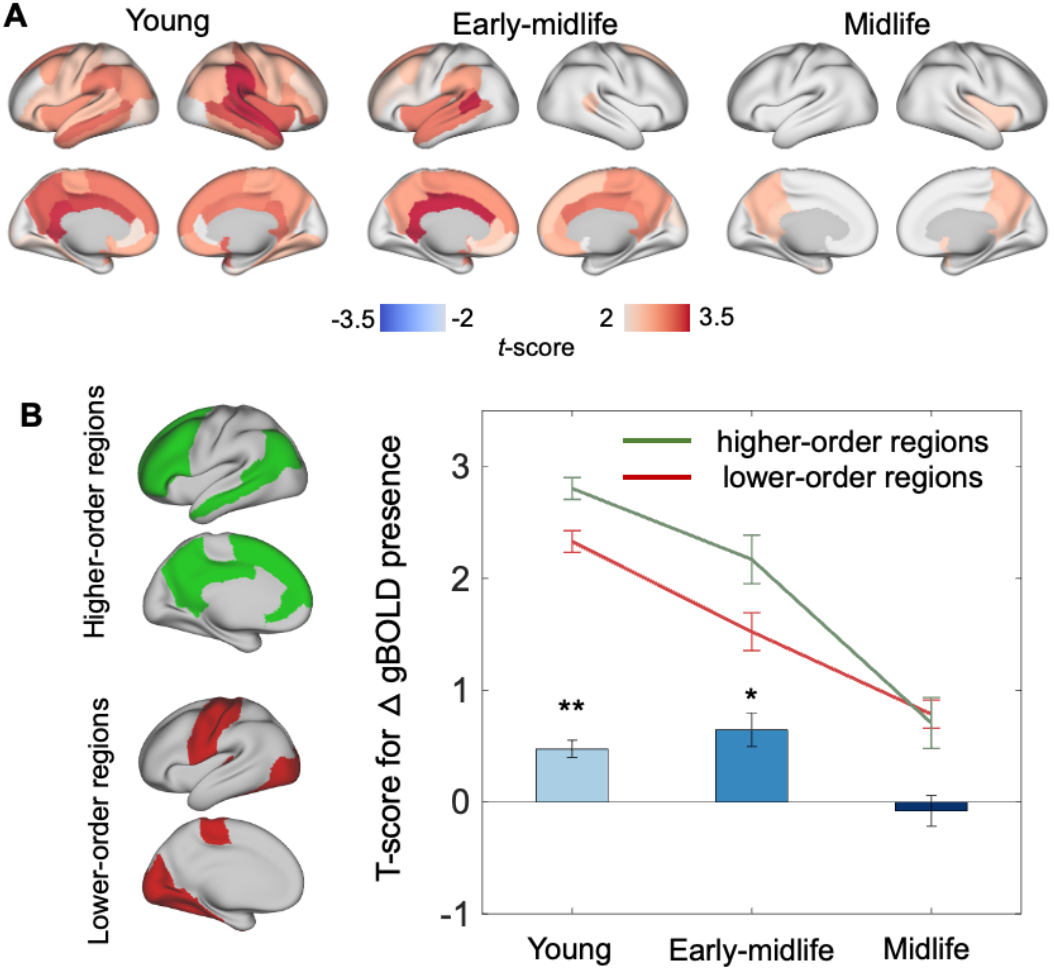
Sleep deprivation-induced changes in gBOLD presence across different age groups. (A)T-score maps showing the TSD-induced changes in gBOLD presence (i.e., the TSD scan minus BS scan) for the three age groups. Brain regions with significant changes (p < 0.05, paired t-test) are color-coded. (B) Summary and comparison of gBOLD presence changes between higher-order cognitive networks (green) and lower-order sensory-motor regions (red). The higher-order and lower-order brain regions are defined based on a brain atlas (left). The t-scores from the maps in (A) are summarized and compared separately for these two sets of brain regions. The higher-order brain regions exhibit a significantly greater increase in gBOLD presence compared to the lower-order brain regions in both the young (p = 0.0014, two-sample t-test) and early-midlife (p = 0.029, two-sample t-test) groups, but no significant differences are observed in the midlife group (p = 0.78, two sample t-test).

## Discussion

Here we demonstrated the age-dependent effects of TSD on infra-slow (<0.1 Hz) global brain activity, which has been linked to AD pathologies, particularly the accumulation of toxic proteins. Specifically, we observed that the global brain activity, measured via gBOLD using fMRI, showed increased amplitude and enhanced coupling with CSF flow after TSD, returning to baseline following recovery sleep. Importantly, this effect, predominantly seen in higher-order brain regions, was present only in individuals aged 20-40 years but not in those aged 40-50 years. Given the growing evidence linking gBOLD activity to brain waste clearance, the TSD-induced increase in gBOLD activity may serve as a compensatory mechanism to counteract impaired clearance due to sleep loss. However, this mechanism appears to be less effective in midlife adults. These findings suggest that age may interact with sleep disturbances to impact brain waste clearance, potentially contributing to the increased risk of dementia in old adults.

We demonstrated that TSD significantly increased the amplitude of gBOLD activity and its coupling with CSF dynamics. The gBOLD signal had been initially considered as non-neuronal noise and was often removed in early resting-state fMRI studies^29,30^, but its high sensitivity to brain arousal state was soon recognized. The fluctuation amplitude of gBOLD increased significantly from wakefulness to light sleep or following the administration of hypnotic drugs^31,32^, whereas decreased after caffeine administration^33^. Recent studies further revealed the neurophysiological basis and spatiotemporal pattern of gBOLD activity. Widespread brain activations at gBOLD peaks are accompanied by specific de-activations in subcortical arousal-regulating nuclei, particularly the basal forebrain nucleus basalis (NB), and a stereotypical electrophysiological pattern indicative of arousal changes^13,34^. Pharmaceutically de-activating the basal forebrain NB on one side of the monkey brain effectively suppressed ipsilateral gBOLD signals^35^. Interestingly, the cortical activations manifest as waves propagating across cortical hierarchies, typically from lower-order sensory-motor areas to higher-order DMN regions, with the early phase accompanied by the de-activation of brainstem nuclei, including the locus coeruleus (LC)^14^. In large-scale neuronal recordings from mice, the infra-slow global brain activity relevant to arousal takes the form of spiking cascades of sequential activations across ∼70% of neuronal populations^15^. Given these early findings, the effect of TSD on gBOLD activity is not entirely surprising and is likely mediated by subcortical neuro-modulatory systems. Our rigorous experimental design ensured that all resting-state fMRI data were collected during wakefulness with subjects’ eyes open, suggesting that large-amplitude gBOLD activity is not exclusive to light sleep^26^. The increased gBOLD amplitude after TSD aligns with the increased global BOLD variability observed previously after partial sleep deprivation^36^. We further showed that TSD also promotes the coupling between gBOLD activity and CSF dynamics.

The TSD-induced increase in the gBOLD-CSF coupling may reflect a compensatory enhancement of clearance function, given recent evidence for its link to toxic protein accumulation in AD. The gBOLD activity has been associated with infra-slow modulations of various physiological signals, such as heart rate and respiratory volume^37–39^. Its coupling with CSF movement, initially found during light sleep, suggests a potential role in CSF-related perivascular clearance^10,11^. It has been hypothesized that the cholinergic and noradrenergic systems modulate at distinct phases of this global dynamic and act synergistically on cerebral arteries to generate vasomotion, thereby facilitating the clearance with the support of simultaneous neural/astrocyte activations^27^. Subsequent studies indeed linked the gBOLD-CSF coupling to various AD pathologies, particularly the accumulation of Aβ and tau^16,17^. More recently, the preferential reduction of gBOLD activity in higher-order DMN regions has been shown to correlate with the early accumulation of Aβ in these regions, thereby linking topological changes in gBOLD with the spreading pattern of Aβ^18^. Based on these findings, the increase in gBOLD activity and gBOLD-CSF coupling after TSD may represent an adaptive increase in clearance function.

This hypothesis of compensatory clearance appears to be consistent with a previous TSD study combining resting-state fMRI and Aβ PET (positron emission tomography) measures. It examined the TSD effects on resting-state BOLD signals in an autonomic network and found a significant increase in the low-frequency (LF, <0.1 Hz) power, which is equivalent to the quantification of the fluctuation amplitude. In TSD subjects, a higher LF power was significantly associated with a lower Aβ burden^40^. While the enhanced clearance function initially appears to contradict previous findings of TSD-induced clearance impairments, this discrepancy may reflect differences in the specific aspects of clearance assessed^6–9^. Prior studies measured the level of Aβ and tau representing a result of clearance processes^7–9^, whereas the gBOLD-CSF coupling likely reflects the instantaneous function of clearance. Thus, the TSD-induced increases in gBOLD activity may represent a compensatory effect that does not fully offset the impaired clearance caused by sleep loss, leading to an overall reduction in waste clearance.

The interaction between age and TSD on gBOLD activity and its coupling with CSF may suggest this compensatory mechanism is impaired in individuals over 40 years old. The previous study observed a similar interaction effect on gBOLD variability, which is however statistically non-significant presumably due to the use of partial sleep deprivation^36^. The total TSD in our study enabled sufficient effect size and confirmed a significant age-TSD interaction effect on gBOLD amplitude, as well as gBOLD-CSF coupling of clearance relevance. Aging is one of the most important risk factors of dementia^25^. Various theories have been proposed to explain the age-related increase in dementia risk^25,41^, and impaired clearance is one of them^42^. Consistent with this hypothesis, the clearance function and gBOLD-CSF coupling have been found to reduce at old ages^22^. The findings of the current study suggest that the age-related reduction in clearance may be partly attributed to the loss of the ability to compensate clearance impairments due to sleep loss. Together with the well-documented sleep disturbances widely observed in older adults, this would highlight a possible pathway through which aging interacts with sleep disturbance to contribute to the age-related risks of dementia. Finally, it is worth noting that the TSD-induced gBOLD changes are more pronounced in the higher-order DMN regions. This topological dominance is similar to those seen at the early stages of Aβ pathology, as well as to the pattern of early Aβ deposition in prodromal/preclinical AD^18^.

## Methods

### Participants and study data

Seventy healthy adults were recruited for a controlled 5-day and 4-night in-laboratory sleep deprivation experiment^26^. Three participants were excluded due to excessive head motion, technical issues, or missing data, leaving 67 participants (aged 34.1 ± 8.9 years, 29 females) for analysis. Among these, 51 participants underwent total sleep deprivation on Night 2, followed by a 12-hour recovery sleep on Night 3 and a subsequent 8-hour recovery sleep on Night 4. The remaining 16 participants served as the control group, maintaining an 8-hour time-in-bed (TIB) schedule across Nights 2–4. To acclimate to the laboratory environment, all participants were provided with a 9-hour TIB sleep opportunity between 9:30 PM and 6:30 AM on Night 1. Resting-state fMRI (rsfMRI) sessions and cognitive tasks were conducted on the mornings of Days 2 (Scan1), 3 (Scan2), and 5 (Scan3) (**Fig. 1A**).

All participants were nonsmokers, right-handed, and confirmed to have no acute or chronic medical or psychological conditions through a comprehensive screening process, which included interviews, medical history reviews, questionnaires, physical examinations, and laboratory tests (blood and urine). The sleep-wake patterns of participants were assessed using multiple methods, including at least 7 days of actigraphy, sleep questionnaires, diaries, and one night of laboratory-based polysomnography and oximetry. All participants maintained consistent sleep routines, with habitual bedtimes between 10:00 PM and 12:00 AM, wake times between 6:00 AM and 9:00 AM, and typical sleep durations of 6.5–8 hours. None of the participants reported habitual napping, irregular sleep schedules, trans-meridian travel, or shift work in the 60 days preceding the study. During the week before and throughout the study, all participants abstained from caffeine, alcohol, tobacco, and medications (except oral contraceptives). Compliance with these requirements and adherence to sleep-wake schedules were verified using actigraphy, sleep diaries, and time-stamped call-ins. The study was approved by the Institutional Review Board of the University of Pennsylvania, adhering to the Declaration of Helsinki, and all participants provided written informed consent. Other aspects of this experiment, which aimed to test various hypotheses, have been previously published^43–48^.

### MR data acquisition and preprocessing

All MR imaging data were acquired using a 3T Tesla MR scanner (Siemens Medical Systems, Erlangen, Germany) with MPRAGE (magnetization-prepared rapid acquisition gradient echo) sequence for high-resolution anatomical imaging. The BOLD fMRI data were collected using a multiband gradient-echo EPI (echo planar imaging) sequence with the following parameters: TR= 2000 milliseconds, TE = 24 milliseconds, FOV = 220*220 mm, matrix size = 64*64*36, and slices thickness = 4 mm. During the resting state scans, participants were instructed to remain still and look at a fixation cross displayed at the center of the screen. All three scans were monitored by an eye-tracker to ensure participants remained awake throughout the procedure.

We conducted preprocessing on the rsfMRI data by employing scripts derived from the 1000 Functional Connectomes Project^49^, slightly customized to suit our needs by using scripts in FSL^50^ and AFNI^51^. The rsfMRI BOLD data were preprocessed, starting with motion correction using the average fMRI image—computed as the mean of all volumes in the time series—as the reference. Subsequent steps included skull stripping, spatial smoothing with a Gaussian kernel (full width at half maximum (FWHM) = 4 mm), temporal filtering to retain frequencies between 0.01–0.1 Hz, and the removal of linear and quadratic temporal trends. Following the registration of the rsfMRI BOLD data to the corresponding high-resolution anatomical images, the data were further aligned to the MNI-152 space. In line with the approach of a prior study^16^, nuisance regression of gBOLD and CSF signals was not performed, as these signals were key variables of interest. Similarly, nuisance regression of head motion parameters was omitted to avoid attenuating the global BOLD signal. While head motion was considered to affect the BOLD signal, recent research has shown that the impact of head motion on the BOLD signal is mediated by transient arousal change^52^.

### gBOLD signal, gBOLD amplitude, CSF signal, and the coupling between the gBOLD signal and CSF signal

The gBOLD signal was calculated by averaging the rsfMRI signals across gray matter regions for each subject (**Fig. 1B & 1C**). A gray matter mask, derived from the Harvard–Oxford cortical and subcortical structural atlas, was used to extract the BOLD signals^53^. To minimize spatial blurring caused by registration, signal extraction was conducted in the original fMRI acquisition space for each subject, with the gray matter mask transformed accordingly. Following extraction, the data were temporally filtered to retain frequencies within the range of 0.01–0.1 Hz. The filtered signals were subsequently normalized by subtracting the mean and dividing by the standard deviation to ensure comparability across subjects. To avoid edge effects from temporal filtering, the first and last five TRs were excluded from the analysis. The gBOLD amplitude was calculated as the standard deviation of the gBOLD signal (**Fig. 1C**).

The CSF signal caused by the inflow effect was extracted from the bottom slices of the rsfMRI data, located near the base of the cerebellum^10,16^ (**Fig. 1B & 1C**). Following the previous study^16^, the rsfMRI data used for CSF extraction only included motion correction, skull stripping, and temporal filtering for preprocessing. The CSF signal for each session was extracted using a manually derived CSF mask from the T2*-weighted fMRI image, which is also validated by the corresponding high-resolution T1-weighted image.

The correlation between gBOLD and CSF signals was computed using the Pearson correlation at different time lags. Similar to previous study^16^, we focused on the correlation at the negative peak (+4 seconds time lag) to represent the strength of the coupling (**Fig. 1D**). We then explored the changes in the strength of coupling after sleep deprivation and recovery sleep, as well as the association of coupling change with age. Paired t-test was conducted to assess the strength of coupling before and after total sleep deprivation within each group. A two-way ANOVA was utilized to explore the interaction between age and scan condition on changes in gBOLD-CSF coupling and gBOLD amplitude.

### gBOLD presence and definition of higher and lower order region

The gBOLD presence signal within specific brain regions was quantified by calculating the correlation between the gBOLD signal and the averaged BOLD signals from individual parcels defined by the DKT-68 parcellation^53^. These correlation values were computed across all parcels, providing a detailed map of gBOLD presence throughout the brain^28^.

To categorize brain regions into lower and higher order, we employed Yeo’s 7-network parcellation framework as used in one previous study^18,54^. Higher-order regions were defined as parcels belonging to the DMN and FPN, both of which are associated with complex cognitive processes (**Fig.3B**). In contrast, lower-order regions were defined as those within the somatomotor and visual networks, which are primarily involved in sensory and motor functions (**Fig.3B)**. This classification facilitated a network-level analysis of gBOLD contributions.

Changes in gBOLD presence were calculated as T-values, representing the differences in gBOLD presence between scans for each parcel. T-values within higher-order and lower-order regions were compared to provide a summary of gBOLD dynamics across distinct functional networks.

### Framewise Displacement

To account for the influence of head movement on our results, we calculated the FD, a metric that quantifies the relative head movement between consecutive time points in the fMRI time series^55^. FD was derived from the six rigid-body motion parameters obtained during motion correction, which included translations along the x, y, and z axes, as well as rotations about these axes. For each time point, the absolute differences in motion parameters relative to the preceding time point were calculated. The FD for each frame was then determined as the sum of these differences. The rotational parameters, initially expressed in degrees, were converted to their equivalent displacement in millimeters using a standard spherical head model with a radius of 50 mm.

We investigated whether head motion influenced the age-specific effects of sleep deprivation on gBOLD-CSF coupling by examining its correlations with both gBOLD-CSF coupling and gBOLD amplitude. To assess the robustness of our findings, we subsequently reanalyzed the primary results after controlling for FD (**Fig.S1 and Fig.S2**).

