## Supplementary Material for "Sleep deprivation exhibits age-dependent effect on infra-slow global brain activity"

**Author Contributions:**

**Y.Y., H.R. &X.L.** contributed to the conception, design of the work, and data analysis; **W.Z., T.M., Y.D., H.R.** contributed to data collection;

**H.R. &X.L.** also devoted the efforts to the supervision, project administration, and funding acquisition;

**Y.Y. &X.L.** contributed to data visualization, and drafting the paper;

**Y.Y., W.Z., Y.M., T.M., Y.D., Y.Y., H.R. &X.L.** contributed to editing and reviewing of the paper;

**Y.Y.** and **W.Z.** are co-first author[†].

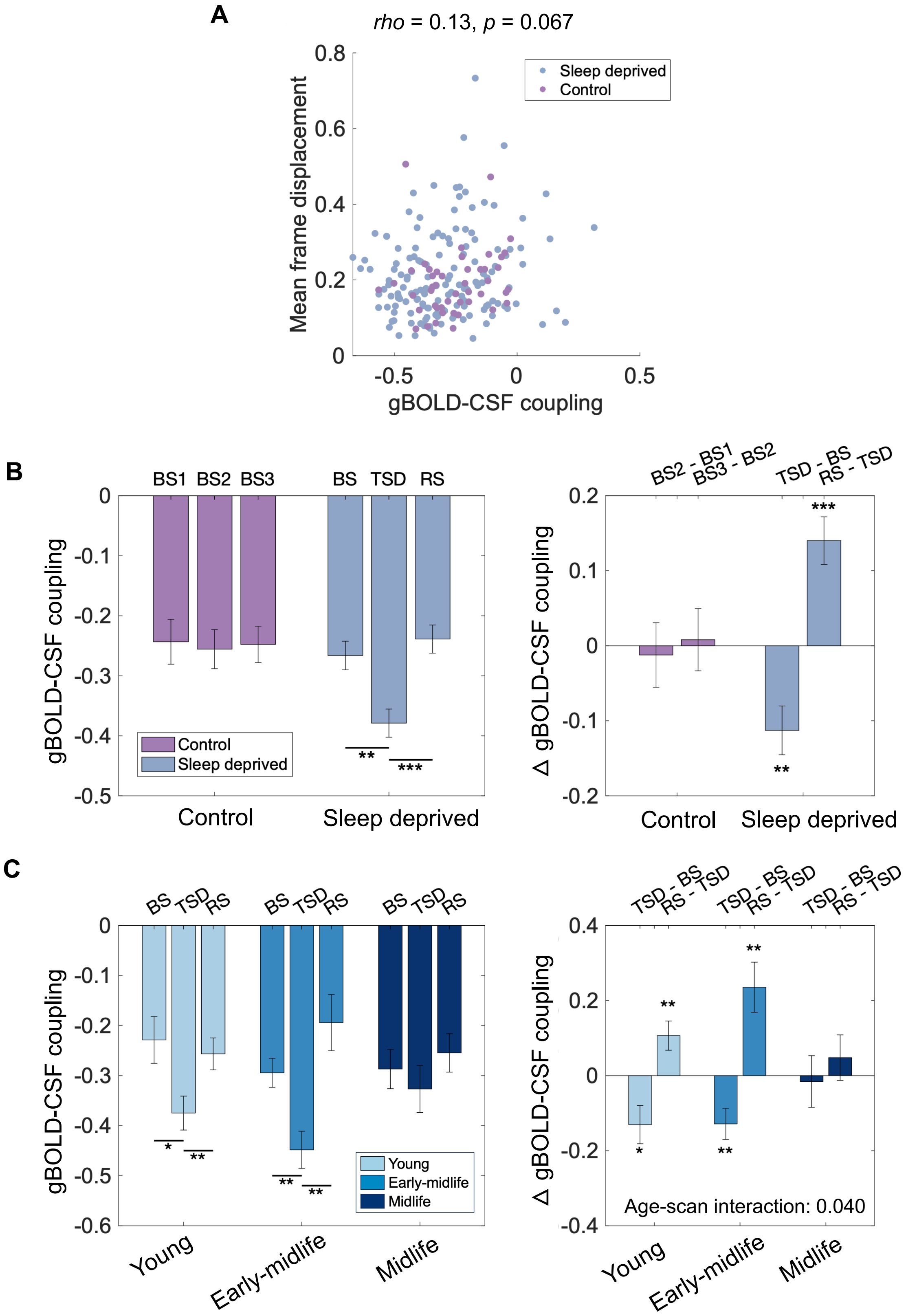
A
rho = 0.13, p = 0.067
Mean frame displacement
gBOLD-CSF coupling
Sleep deprived
Control
B
BS1
BS2
BS3
BS
TSD
RS
gBOLD-CSF coupling
Control
Sleep deprived
BS2 - BS1
BS3 - BS2
TSD - BS
RS - TSD
Δ gBOLD-CSF coupling
C
Young
Early-midlife
Midlife
Age-scan interaction: 0.040

***Figure S1. The impact of TSD on gBOLD-CSF coupling remained robust after regressing out the head motion effect. (A)*** *The head motion showed no significant correlation with gBOLD-CSF coupling (rho = 0.13, p = 0.067, Spearman's correlation).* ***(B)*** *After adjusting for the head motion parameter, the gBOLD-CSF coupling (left) and its changes (right) over the three scans in sleep-deprived and control groups. Subjects who underwent TSD showed a significant increase in gBOLD-CSF coupling during TSD scan compared to BS scan (p = 0.0011, paired t-test), and the coupling returned to baseline during the RS scan (p = 0.000053, paired t-test). The gBOLD-CSF coupling showed no significant changes (BS2 vs. BS1, p = 0.78; BS3 vs. BS2, p = 0.85, paired t-test) over the three scans in the control group. (**C**) After adjusting for head motion parameter, the changes in coupling across scans were significant for both young (TSD vs. BS, p = 0.012; RS vs. TSD, p = 0.0045, paired t-test) and early-midlife (TSD vs. BS, p = 0.0026; RS vs. TSD, p = 0.0020, paired t-test) groups, but not for the midlife group (TSD vs. BS, p = 0.56; RS vs. TSD, p = 0.25, paired t-test). Age and scan showed a significant interaction effect on changes in gBOLD-CSF coupling (p = 0.040, two-way ANOVA). Error bars represent the standard error of the mean (SEM) across subjects. Asterisks represent the significance level (*: 0.01 < p < 0.05; **: 0.001 < p < 0.01; ***p < 0.001)*

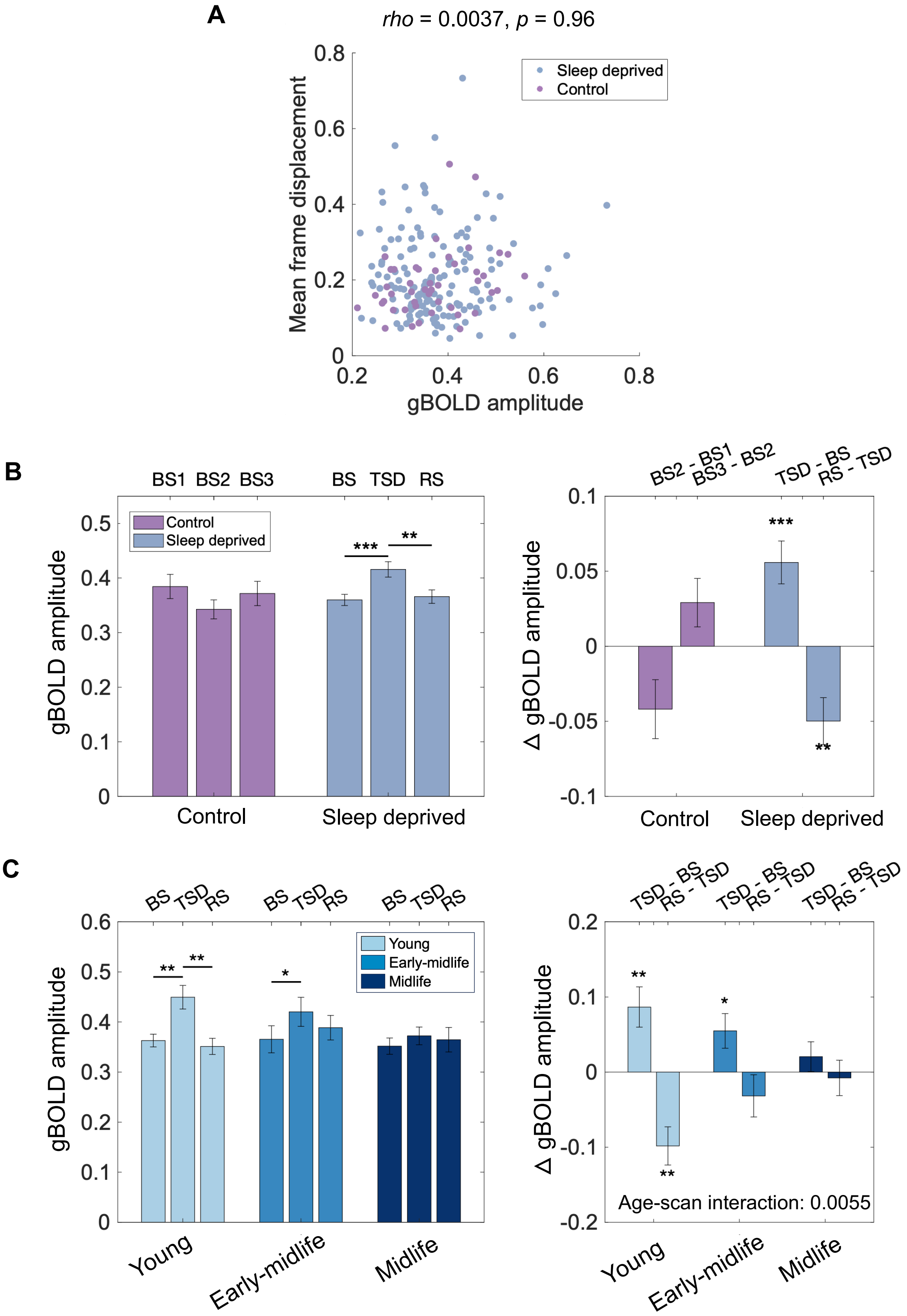
A
rho = 0.0037, p = 0.96
Sleep deprived
Control
Mean frame displacement
gBOLD amplitude
0
0.2
0.4
0.6
0.8
B
BS1 BS2 BS3
BS TSD RS
Control
Sleep deprived
***
**
gBOLD amplitude
0
0.1
0.2
0.3
0.4
0.5
BS2 - BS1
BS3 - BS2
TSD - BS
RS - TSD
Δ gBOLD amplitude
0.1
0.05
-0.05
-0.1
C
BS TSD RS
Young
Early-midlife
Midlife
*
0.6
TSD - BS
RS - TSD
0.2
-0.2
Age-scan interaction: 0.0055

***Figure S2. The impact of TSD on gBOLD amplitude remained robust after regressing out the head motion effect. (A)*** *The head motion showed no significant correlation with gBOLD amplitude (rho = 0.0037, p = 0.96, Spearman's correlation).* ***(B)*** *After adjusting for the head motion parameter, the gBOLD amplitude (left) and its changes (right) over the three scans in sleep-deprived and control groups. Subjects who underwent TSD showed a significant increase in gBOLD amplitude during TSD scan compared to BS scan (p = 0.00026, paired t-test), and it returned to baseline in the RS scan (p = 0.0024, paired t-test). The gBOLD amplitude showed no significant changes (BS2 vs. BS1, p = 0.05; BS3 vs. BS2, p = 0.09, paired t-test) over the three scans in the control group. (**C**) After adjusting for head motion parameter, the changes in gBOLD amplitude after sleep deprivation across scans were significant for both young (TSD vs. BS, p = 0.0044, paired t-test) and early-midlife (TSD vs. BS, p = 0.033, paired t-test) groups. Only for young group, the gBOLD amplitude returned to baseline level after recovery sleeps (RS vs. TSD, p = 0.0010, paired t-test), but not in early-midlife group (RS vs. TSD, p = 0.28, paired t-test). No significant changes were observed in midlife group (TSD vs. BS, p = 0.32; RS vs. TSD, p = 0.74, paired t-test). Age and scan showed a significant interaction effect on changes in gBOLD amplitude (p = 0.0055, two-way ANOVA).*

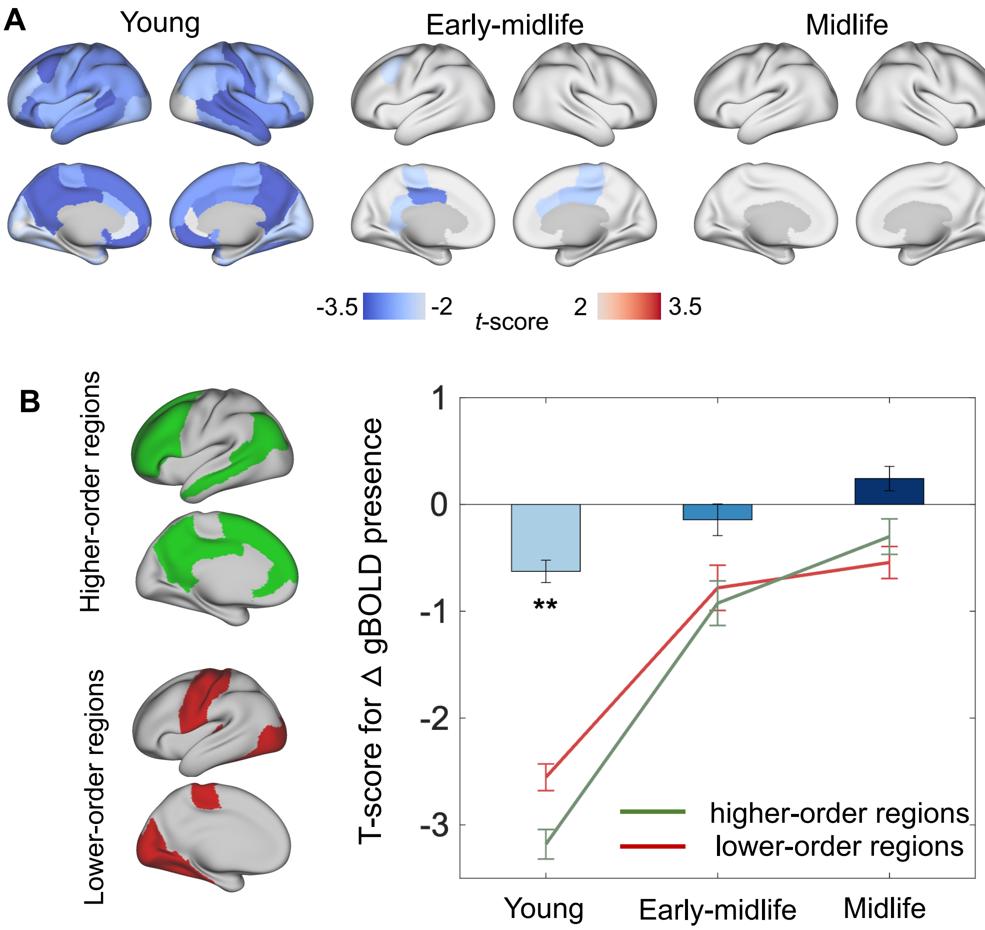


***Figure S3. The changes in gBOLD presence across different age groups after recovery sleeps.*** ***(A)*** *T-score maps showing the gBOLD presence change after recovery sleep (i.e., the RS scan minus TSD scan) for the three age groups. The brain regions with significant changes ($p < 0.05$, paired t-test) are color coded.* ***(B)*** *Summary and comparison of gBOLD presence changes between higher-order cognitive networks (green) and lower-order sensory-motor regions (red). The higher-order and lower-order brain regions were defined based on a brain atlas (left). The t-scores from the maps in* ***(A)*** *were summarized and compared separately for these two sets of brain regions. The higher-order brain regions exhibited a significantly greater decrease in gBOLD presence compared to lower-order brain regions in young group ($p = 0.0020$, two-sample t-test), but not the early-midlife ($p = 0.64$, two-sample t-test)and midlife ($p = 0.30$, two-sample t-test) groups.*
